# A program to automate the discovery of drugs for West Nile and Dengue virus – programmatic screening of over a billion compounds on PubChem, generation of drug leads and automated *In Silico* modelling

**DOI:** 10.1101/2020.06.17.148312

**Authors:** A S Ben Geoffrey, Akhil Sanker, Rafal Madaj, Mario Sergio Valdés Tresanco, Manish Upadhyay, Judith Gracia

## Abstract

Our work is composed of a python program for programmatic data mining of PubChem to collect data to implement a machine learning based AutoQSAR algorithm to generate drug leads for the flaviviruses – Dengue and West Nile. The drug leads generated by the program are feed as programmatic inputs to AutoDock Vina package for automated *In Silico* modelling of interaction between the compounds generated as drug leads by the program and the chosen Dengue and West Nile drug target methyltransferase, whose inhibition leads to the control of viral replication. The machine learning based AutoQSAR algorithm involves feature selection, QSAR modelling, validation and prediction. The drug leads generated each time the program is run is reflective of the constantly growing PubChem database is an important dynamic feature of the program which facilitates fast and dynamic drug lead generation against the West Nile and Dengue virus in way which is reflective of the constantly growing PubChem database. The program prints out the top drug leads after screening PubChem library which is over a billion compounds. The leads generated by the program are fed as programmatic inputs to an *In Silico* modelling package. The interaction of top drug lead compounds generated by the program and drug targets of West Nile and Dengue virus, was modelled in an automated way through programmatic commands. Thus our program ushers in a new age of automatic ease in the virtual drug screening and drug identification through programmatic data mining of chemical data libraries and drug lead generation through machine learning based AutoQSAR algorithm and an automated *In Silico* modelling run through the program to study the interaction between the drug lead compounds and the drug target protein of West Nile and Dengue virus

## Introduction

PubChem database is a publically accessible bio and chemical data repository of over a billion compounds and their bioactivity data [1]. The programmatic access to PubChem database can be accomplished through web API packages such as PUB-REST and PyPubChem by which data mining for a particular task can be automated by programmatic access to the database using python commands [2, 3]. Statistically predicted drug leads can be generated from a QSAR model for a particular pharmacological activity [4–7]. However the data sets used in QSAR modelling has to be curated each time by researchers’ before carrying out the study. In our present efforts, we have automated the process of data set fetching required for a QSAR study which is associated with a particular drug target by programmatic access to PubChem database through python commands. Therefore, each time the program is run, the drug leads generated each time are reflective of the constantly growing PubChem database. Programmatic data mining was carried out on PubChem to collect data to implement a machine learning based AutoQSAR algorithm to generate drug leads for the flaviviruses – Dengue and West Nile. The drug leads generated by the program are feed as programmatic inputs to AutoDock Vina package for automated *In Silico* modelling of interaction between the drug lead compounds and the chosen Dengue and West Nile drug target methyltransferase [8,9], whose inhibition leads to the control of viral replication. Thus our program ushers in a new age of programmatic ease in the virtual drug screening and identification of drugs against the West Nile virus and Dengue virus through programmatic data mining of chemical data libraries and drug lead generation through machine learning based AutoQSAR algorithm and an automated In Silico analysis of the drug lead compounds.

While previous such attempts [10] do not use programmatic virtual screening, the ones that use programmatic screening [11, 12] use pre-downloaded data sets from chemical libraries. We make a case for the uniqueness of our work in the fact that while we use programmatic screening, the data set fetching happens in real time when the program is run and thus the results are reflective of constantly growing chemical data libraries. We also make a case for the uniqueness of our work in that fact that among similar attempts at programmatic data mining and drug lead generation through machine learning based AutoQSAR algorithms [12–14], our code implements automated *In Silico* modelling by coupling the drug leads generated by the AutoQSAR algorithm as programmatic inputs to *In Silico* modelling packages. While the work brings to published literature novelty in methods and techniques employed, it also brings new scientific findings to published literature as the West Nile and Dengue virus drug target methyltransferase, has not been approached by data-driven machine learning based drug discovery methods. Thus, in this way the entire process of virtual screening has been automated and our program package can be used as a complete automated virtual screening suite to identify West Nile and Dengue virus drugs in ways unique ways as detailed above as compared to existing works. The drug leads identified by the program can further be carried forward to *In Vitro* and *In Vivo* experimental testing

## Methods and Techniques

The workflow of the algorithm implemented as code is shown in Fig.1. The first process involved is programmatic data mining the PubChem database to automate the process of data acquisition to implement a machine learning based AutoQSAR algorithm to automate the process of generation of drug leads against Flaviviruses – West Nile and Dengue virus. This is accomplished through programmatic access to PubChem database through python commands [15, 16]. The second process involves implementing a machine learning based AutoQSAR algorithm. The workflow of a machine learning based AutoQSAR involves feature learning and descriptor selection, QSAR modelling, validation and prediction [17–19]. The next process involves a programmatic querying of PubChem database to generate drug leads from compounds in PubChem that are structurally associative to compounds known to be active against the Flavivirus drug target. The drug leads are also required to satisfy the Lipinski’s criteria to qualify as an orally active drug in humans [20].

**Fig 1.**
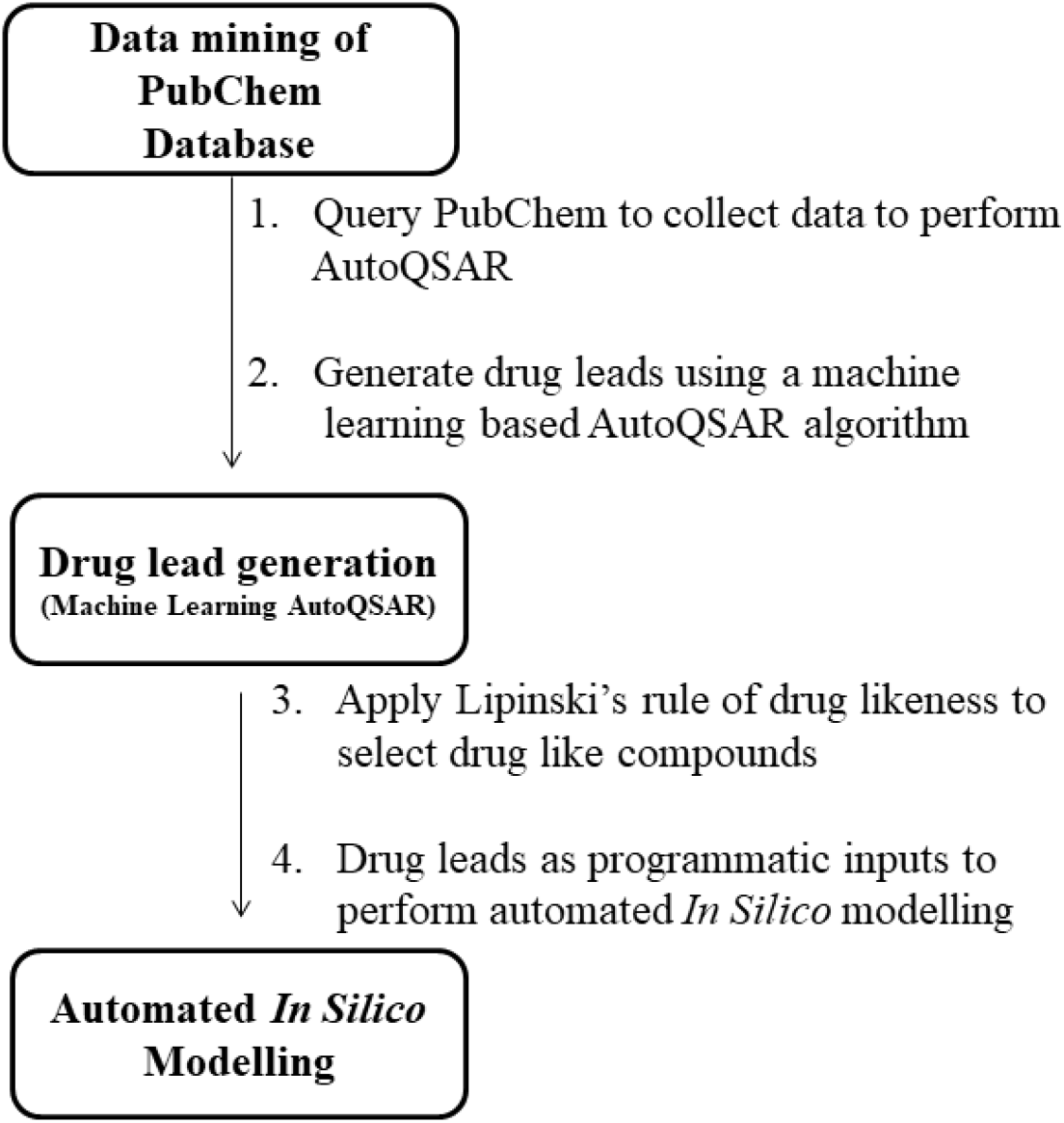
Block diagrammatic representation of the workflow of the algorithm

Running the program requires no more programming knowledge than running the python executable file in python 3 environment in Linux OS along with some python dependency packages installed such as:

pandas==1.0.1
numpy
matplotlib
scikit-learn
seaborn

Other additional packages required to perform automated virtual screening

openbabel 2.4.1
mgltools 1.5.4
autodock-vina 1.1.2-4

The program is hosted, maintained and supported at the GitHub repository link given below https://github.com/bengeof/Automation-in-the-discovery-of-drugs-for-Flaviviruses---West-Nile-and-Dengue-virus

The run time of the program is expected to be a few hours with variation based on CPU and internet speed. As final output the program prints out the top 30 drug leads which are identified with PubChem CIDs that are printed out and the final output is shown in Fig.2

**Fig.2.**
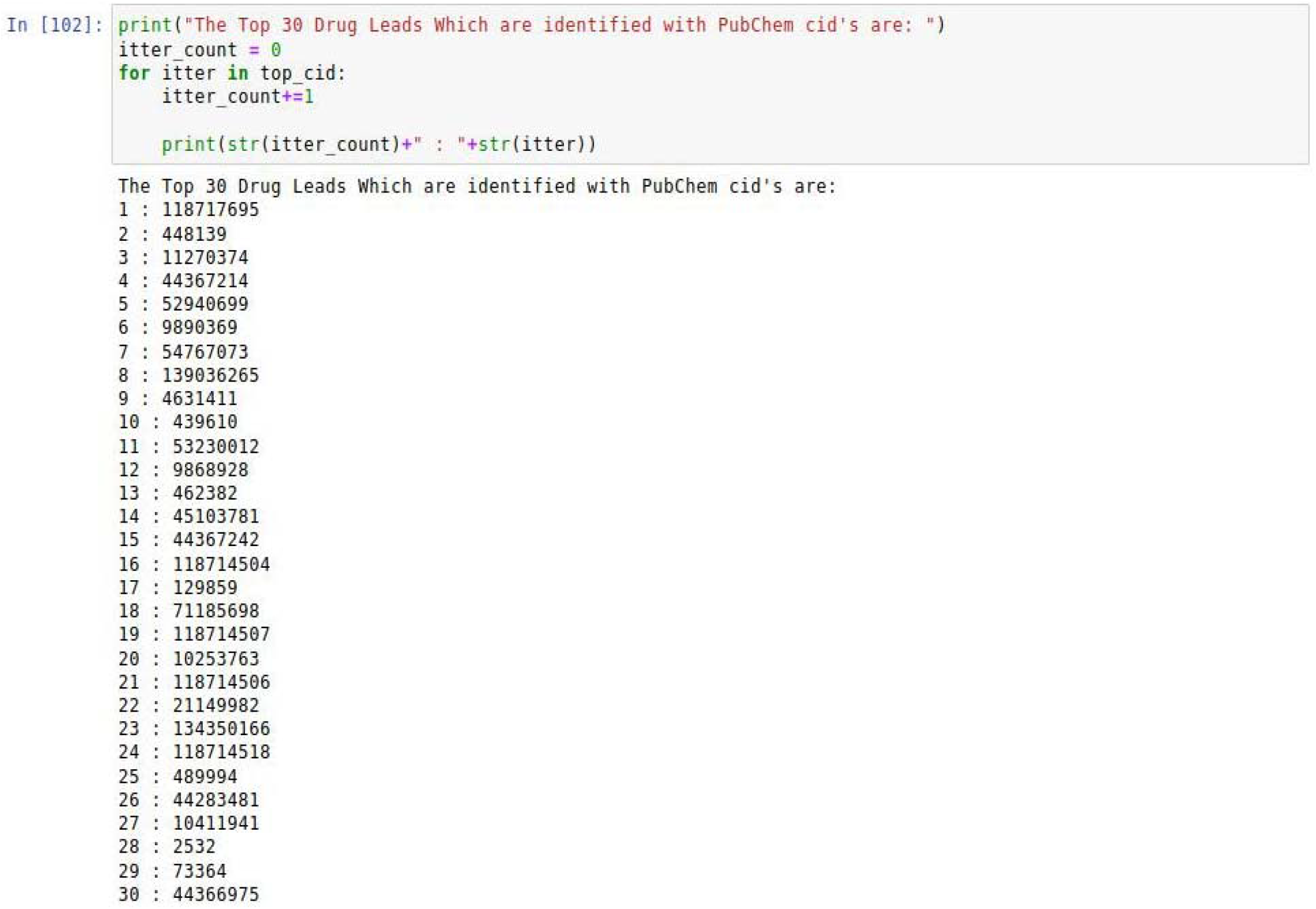
Drug leads identified with PubChem CIDs generated as the output of the program

The crystal structure of the West Nile virus drug target was downloaded from the RCSB-PDB database [21] with PDB ID : 2Oy0. The crystal structure of the Dengue virus drug target was downloaded from the RCSB-PDB database [21] with PDB ID : 1L9K. The pdbqt files of drug target proteins prepared for automated docking were kept in the working folder. The structure of the drug leads generated by the program were programmatically downloaded from PubChem and programmatically converted to pdbqt format through OpenBabel in the program. The ligand files were also programmatically prepared for docking through use of AutoDock scripts for ligand preparation. The molecular docking process of AutoDock Vina [22, 23] was initiated programmatically through the program and the interaction between the drug targets proteins and lead drug compounds were studied. The program allows the user to control the number of lead compounds that must proceed for *In Silico* analysis in the way as shown in Fig.3. The results were visualized in Discovery studio [24]

**Fig. 3.**
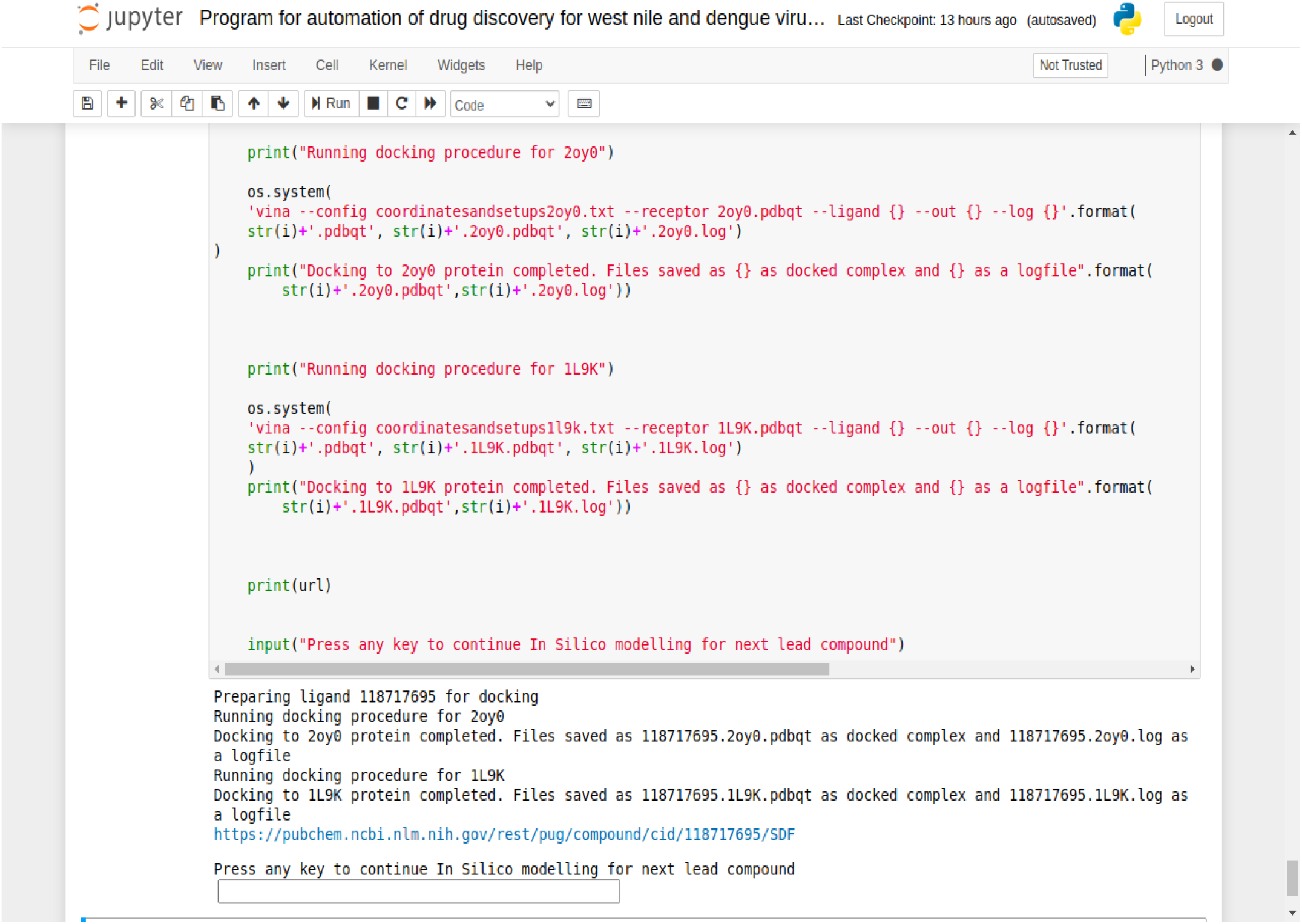
Output screen of the automated *In Silico* modelling

## Results and Discussion

The python executable file was run in Python 3 environment with the dependency packages mentioned above. The run time of the program is about a few hours however it is expected to vary depending on CPU and internet speed on different machines. The output screen of the program is shown in Fig.2. The program prints out the PubChem CID of compounds that are the best 30 possible drug leads against Flaviviruses. It does so by an automated programmatic data mining of PubChem database and by implementing a machine learning based AutoQSAR algorithm on the dataset to generate the drug leads. The drug leads generated by the program for the West Nile and Dengue virus are useful to screen PubChem database which is over a billion compounds and generate drug leads for West Nile and Dengue virus that are useful to further pursue *In Silico, In Vitro and In Vivo* testing and is expected to save computational and experimental testing costs for the pharmaceutical industry.

The drug leads generated by the program were required to fit the Lipinski’s criteria of drug likeness. Compounds satisfying the Lipinski’s are likely to be orally active drugs in humans. The structure of the drug lead compounds were programmatically downloaded from PubChem through the program and they were fed as programmatic inputs to a *In Silico* modelling package used widely, AutoDock-Vina. Therefore the study of the interaction of the drug lead compounds and the West Nile virus drug target methyltransferase (PDB ID :2oy0) and the Dengue virus drug target methyltransferase (PDB ID :1L9K) was automated through the programmatic inputs given to AutoDock-Vina in the program. The results of the molecular docking of the lead compounds and West Nile virus drug target methyltransferase were found to possess favourable interaction between the lead compounds and the target methyltransferase, thus making a case for the usefulness of our tool in discovering drugs for West Nile virus. Select results are given in Table 1. The interaction of the lead compounds generated by the program and the West Nile virus drug target in shown in Fig.4a, 4b & 4c. The compound identified with the PubChem CID 439610 formed a stable complex with the West Nile virus drug target with a binding energy of −7.4 and the interacting residues were found to be Glu218, Gly81, Lys182, Asp146, Glu111, Cys82, Ser56, Gly58. Similarly the compounds generated as drug leads by the program also interacted favourably with the drug target of Dengue virus and select results are tabulated in Table 2 and the interaction between the compounds the target protein in shown in Fig.5a, 5b & 5c. It was found that the compound identified with the PubChem CID 439610 formed a stable complex with the Dengue virus drug target with a binding energy of −8.1 and the interacting residues were found to be Asp79, Gly81, Glu111, SO4:903, Lys181, Lys61, Asp146. Thus again making a case for the usefulness of the programmatic tool in identifying drugs against Dengue virus.

**Fig.4a.**
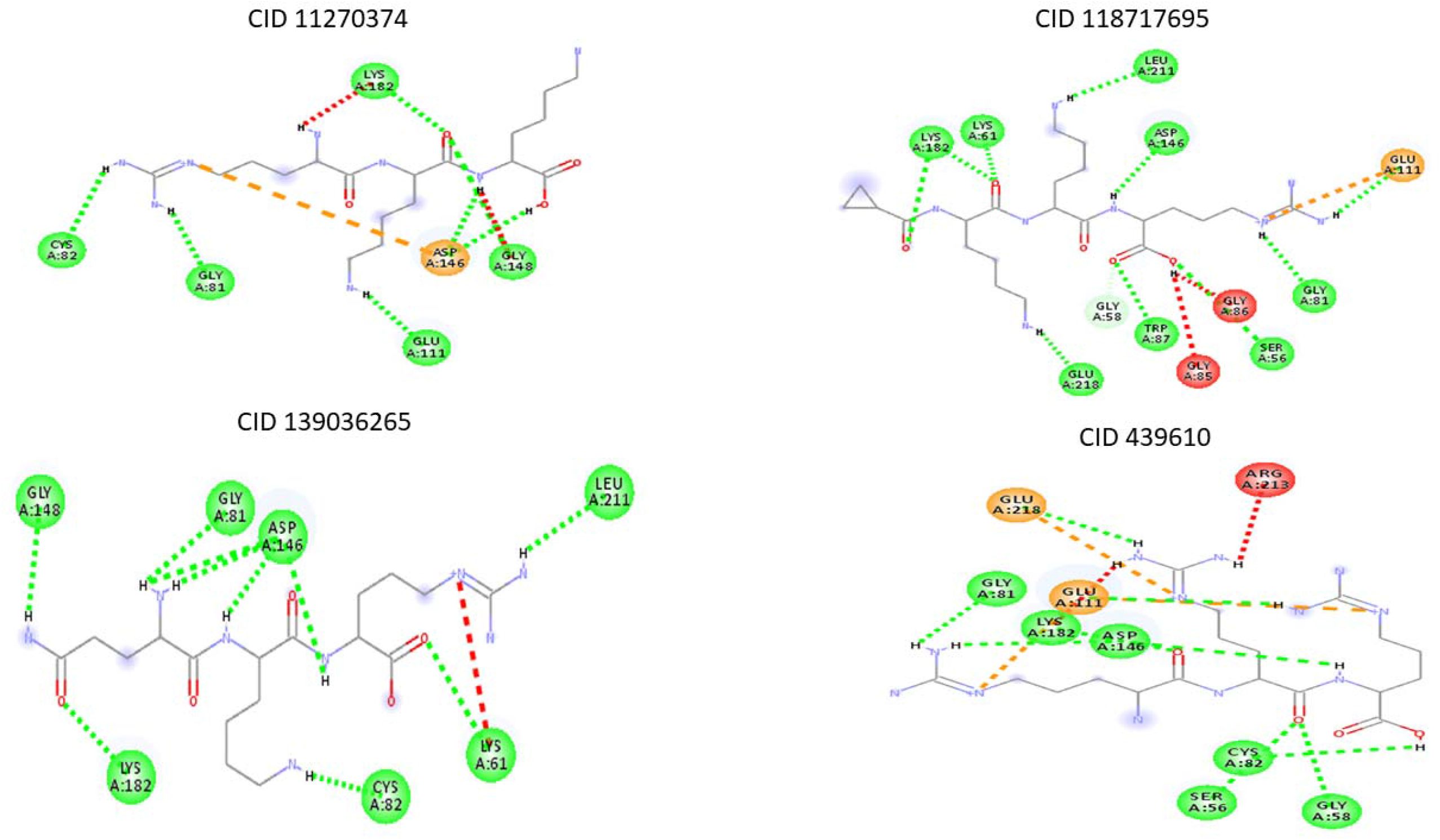
Interaction of drug leads compounds generated by the program and the West Nile Virus drug target protein

**Fig.4b.**
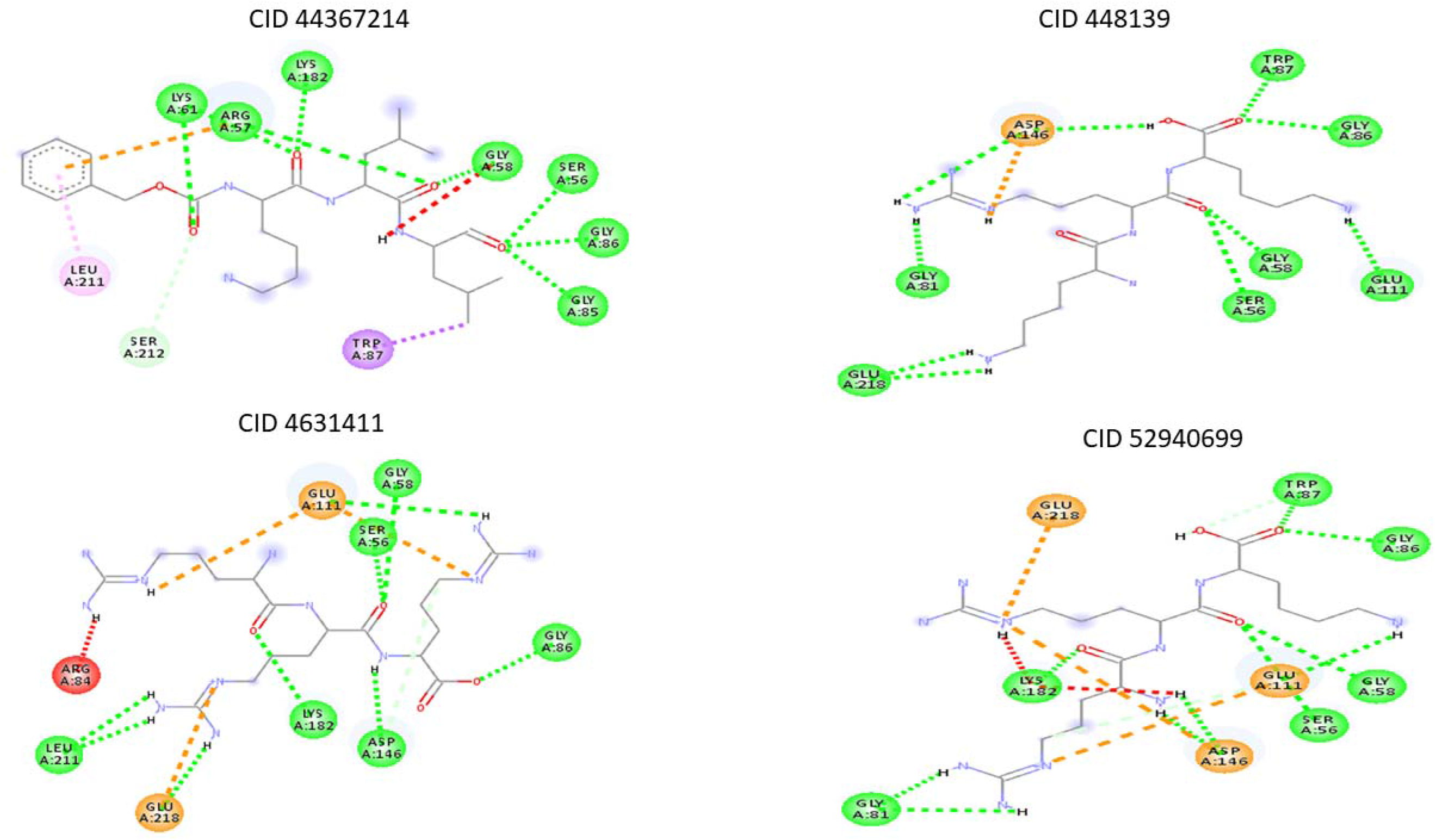
Interaction of drug leads compounds generated by the program and the West Nile Virus drug target protein

**Fig.4c.**
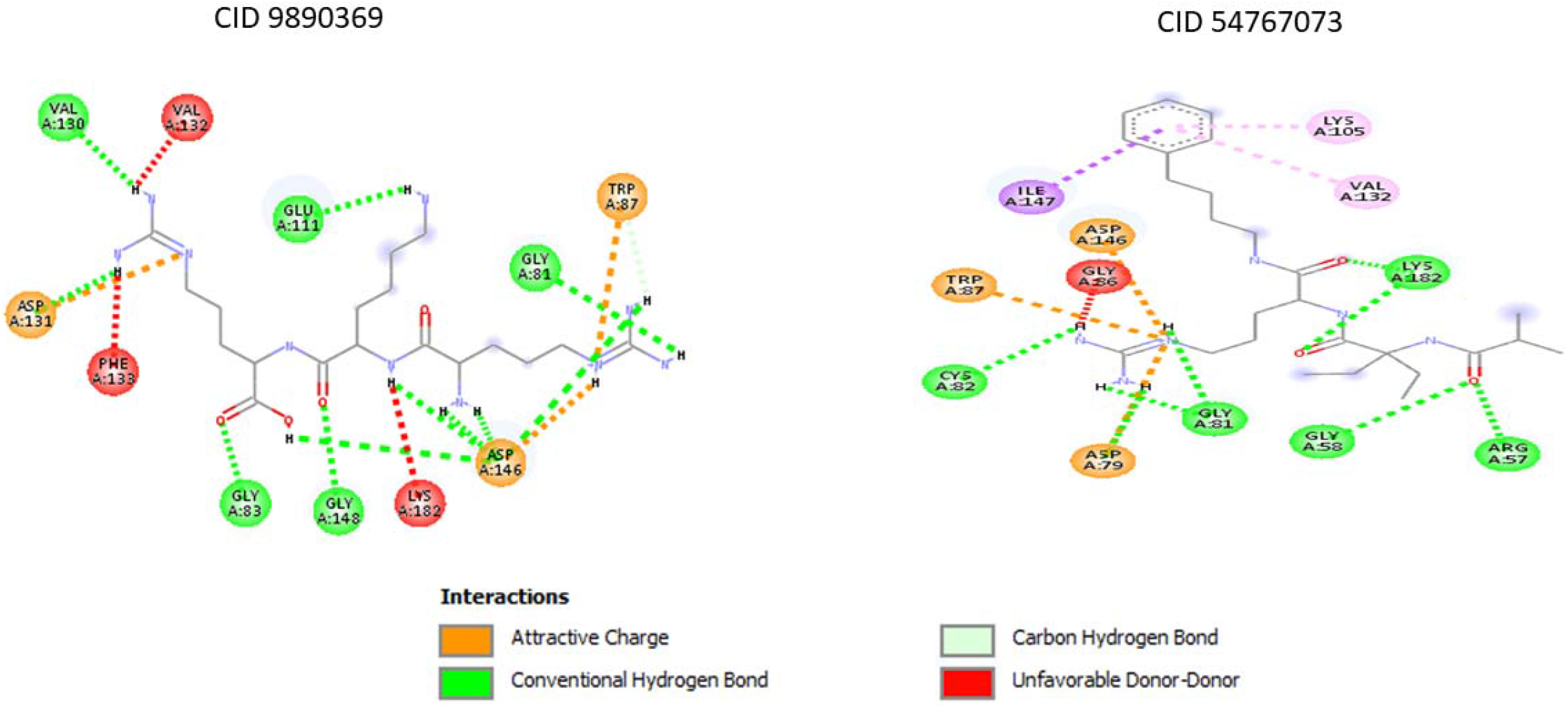
Interaction of drug leads compounds generated by the program and the West Nile Virus drug target protein

**Fig.5.**
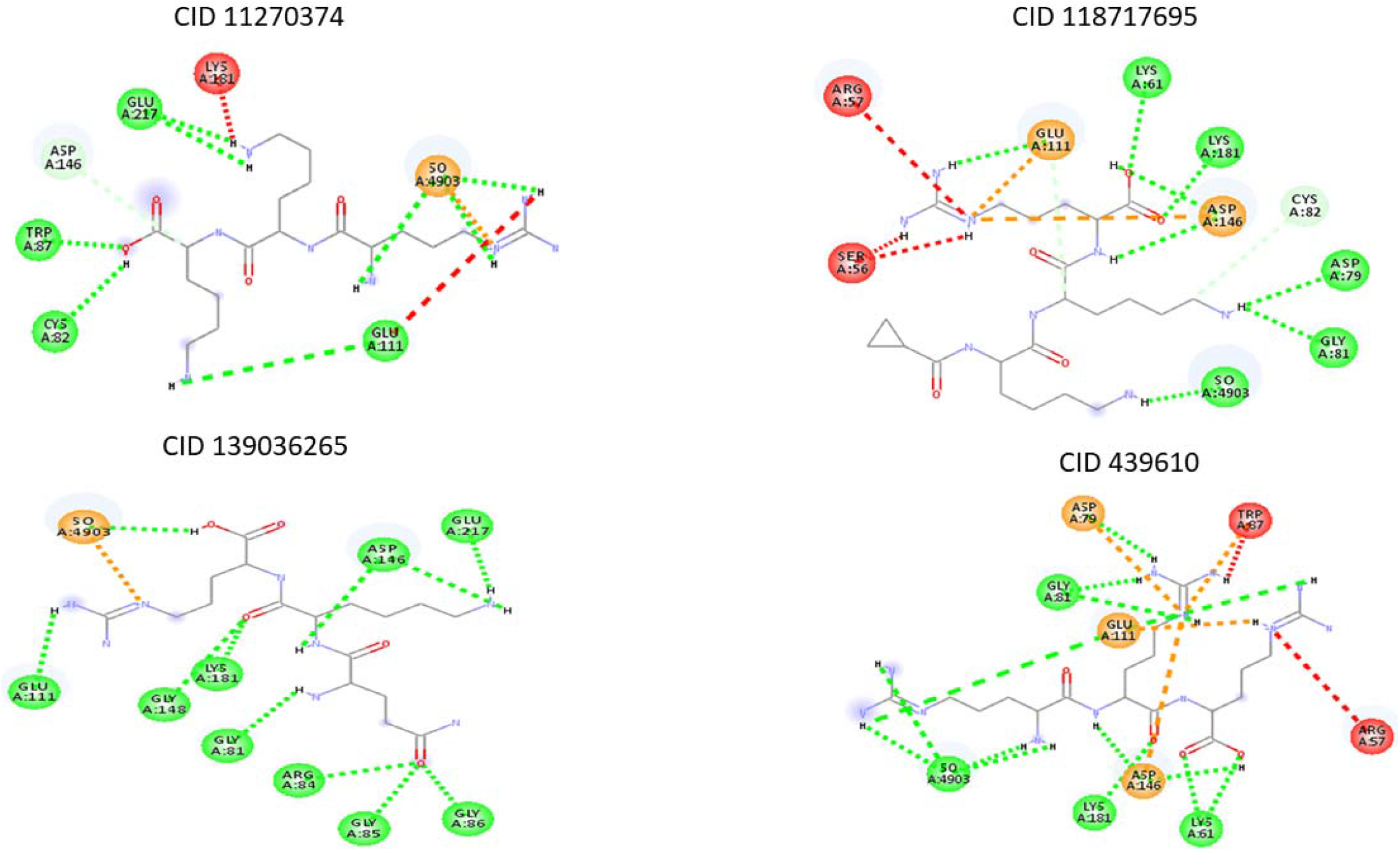
Interaction of drug leads compounds generated by the program and the Dengue virus drug target protein

**Fig.5b.**
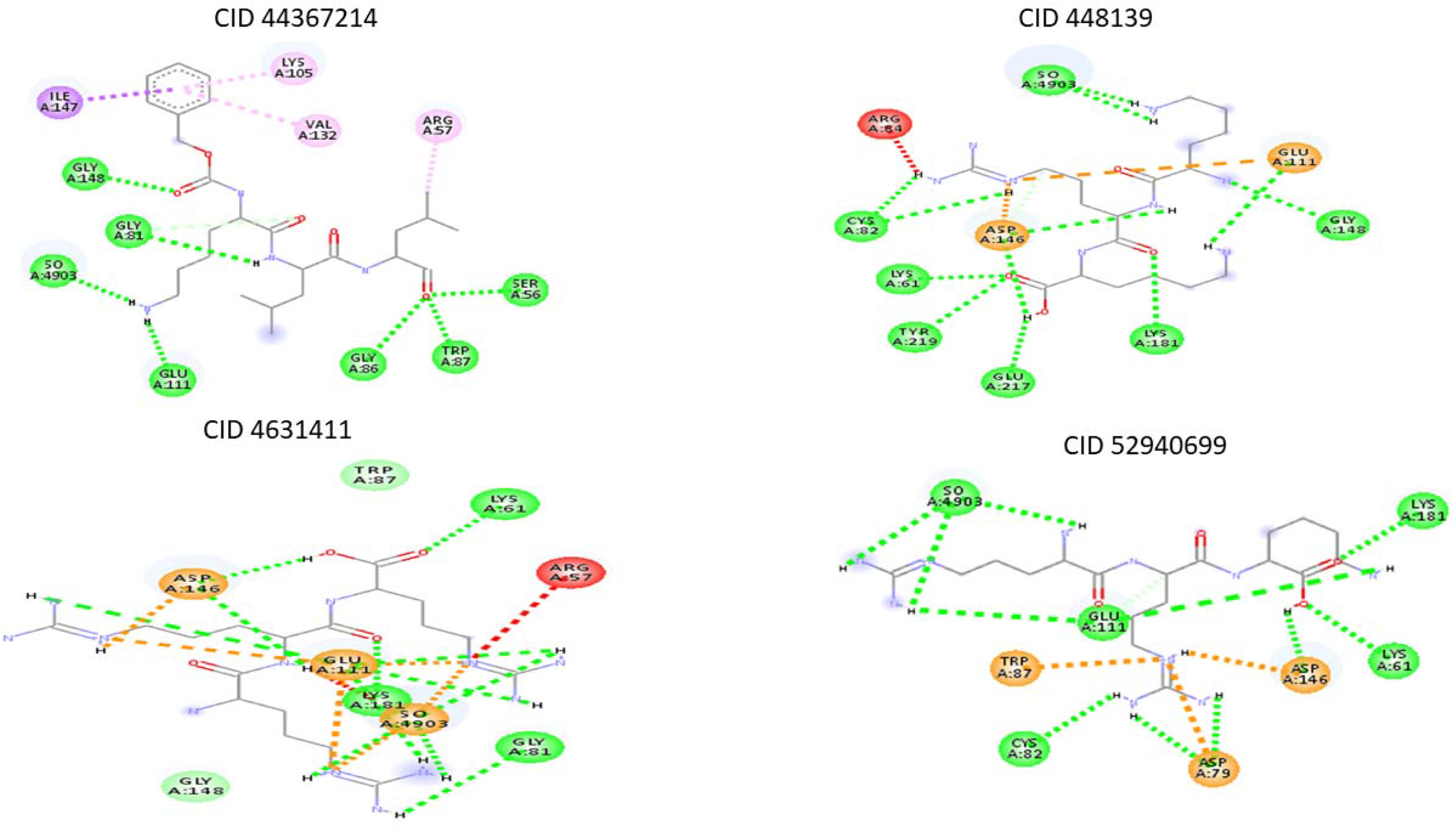
Interaction of drug leads compounds generated by the program and the Dengue virus drug target protein

**Fig.5c.**
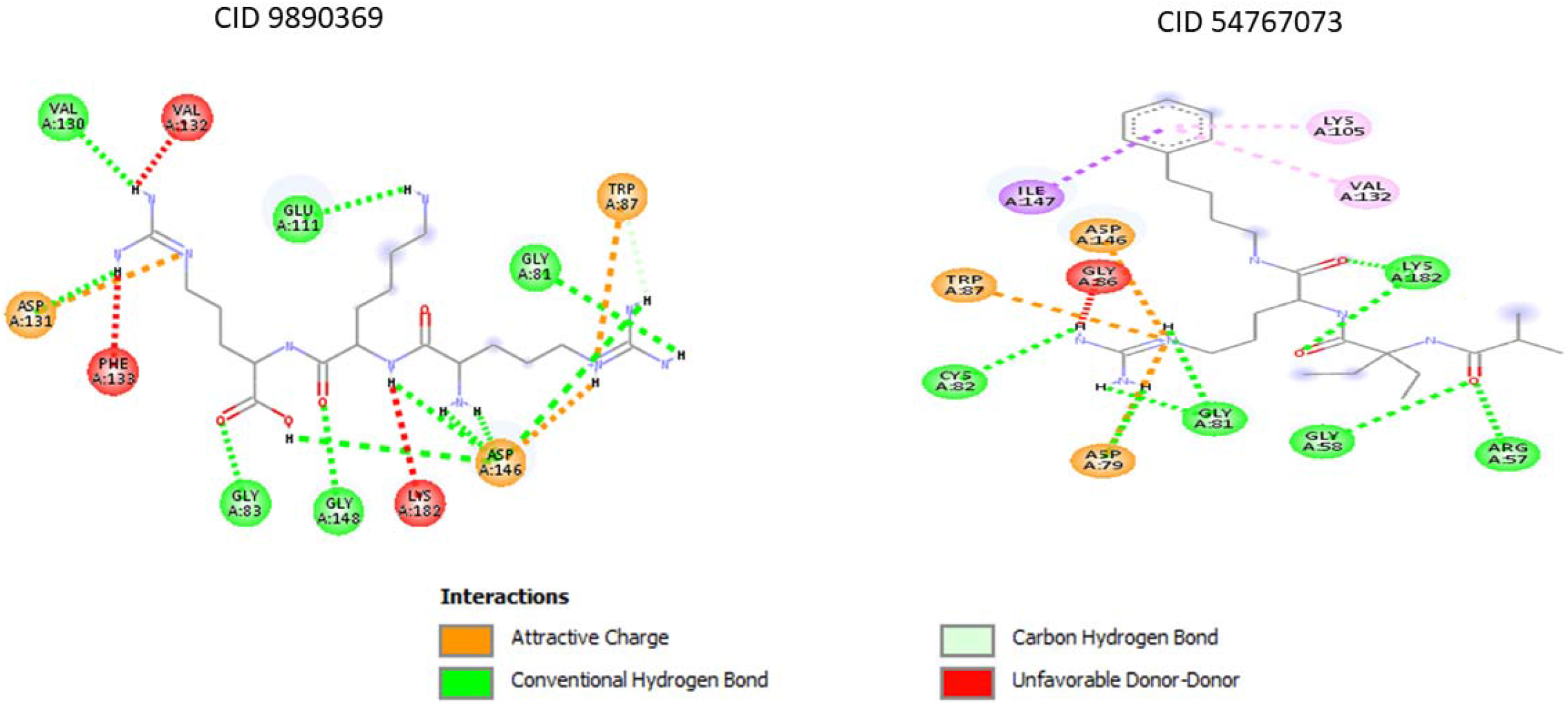
Interaction of drug leads compounds generated by the program and the Dengue virus drug target protein

**Table 1.** Interaction of select drug lead compounds and the drug target protein of West Nile virus

| Ligand | Target protein | Docking score | Hydrogen bonded residues<br>A Chain |
| --- | --- | --- | --- |
| CID118717695 | 2oY0 | -7.1 | Gly81, **Ser56**, Trp87, Glu218, **Asp146**, Leu211, Lys61, **Glu111**, Lys182 |
| CID448139 |  | -6.9 | **Glu111**, Gly58, **Ser56**, Glu218, Gly81, **Asp146**, Trp87, Gly86 |
| CID11270374 |  | -6.8 | **Asp146**, Gly148, **Glu111**, Gly81, **Cys82**, Lys182 |
| CID44367214 |  | -7.4 | Gly85, Gly86, **Ser56**, Gly58, Lys182, Arg57, Lys61 |
| CID52940699 |  | -6.8 | Trp87, Gly86, Gly58, **Ser56**, **Glu111**, **Asp146**, Lys182, Gly81 |
| CID9890369 |  | -6.9 | Gly81, **Asp146**, Gly83, Gly148, Asp131, Val130, **Glu111** |
| CID54767073 |  | -7.6 | Arg57, Gly58, Gly81, **Cys82**, Lys182 |
| CID139036265 |  | -6.7 | Leu211, Lys61, **Cys82**, Lys182, Gly148, Gly81, **Asp146** |
| CID4631411 |  | -7.4 | **Ser56**, Gly58, **Glu111**, Gly86, **Asp146**, Lys182, Glu218, Leu211 |
| CID439610 |  | -7.4 | Glu218, Gly81, Lys182, **Asp146**, **Glu111**, **Cys82**, **Ser56**, Gly58 |

**Table 2.**
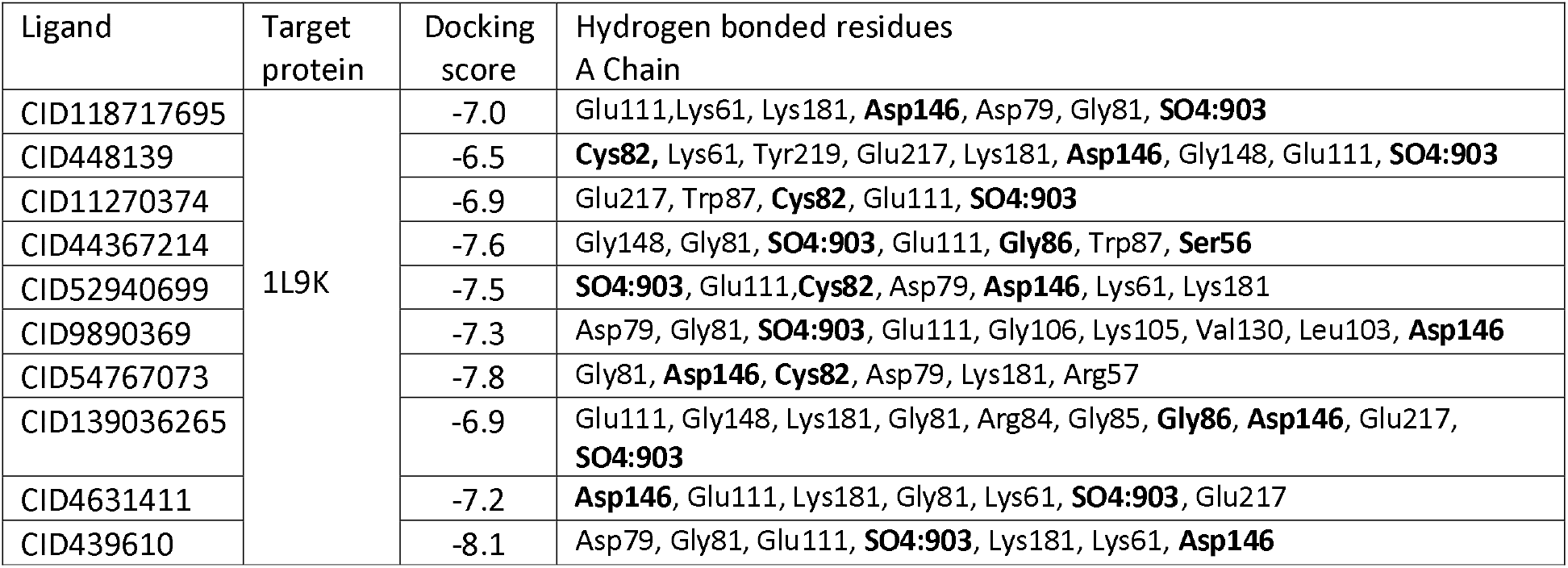
Interaction of select drug lead compounds and the drug target protein of Dengue virus

| Ligand | Target protein | Docking score | Hydrogen bonded residues<br>A Chain |
| --- | --- | --- | --- |
| CID118717695 | 1L9K | -7.0 | Glu111, Lys61, Lys181, **Asp146**, Asp79, Gly81, **SO4:903** |
| CID448139 |  | -6.5 | **Cys82**, Lys61, Tyr219, Glu217, Lys181, **Asp146**, Gly148, Glu111, **SO4:903** |
| CID11270374 |  | -6.9 | Glu217, Trp87, **Cys82**, Glu111, **SO4:903** |
| CID44367214 |  | -7.6 | Gly148, Gly81, **SO4:903**, Glu111, **Gly86**, Trp87, **Ser56** |
| CID52940699 |  | -7.5 | **SO4:903**, Glu111, **Cys82**, Asp79, **Asp146**, Lys61, Lys181 |
| CID9890369 |  | -7.3 | Asp79, Gly81, **SO4:903**, Glu111, Gly106, Lys105, Val130, Leu103, **Asp146** |
| CID54767073 |  | -7.8 | Gly81, **Asp146**, **Cys82**, Asp79, Lys181, Arg57 |
| CID139036265 |  | -6.9 | Glu111, Gly148, Lys181, Gly81, Arg84, Gly85, **Gly86**, **Asp146**, Glu217, **SO4:903** |
| CID4631411 |  | -7.2 | **Asp146**, Glu111, Lys181, Gly81, Lys61, **SO4:903**, Glu217 |
| CID439610 |  | -8.1 | Asp79, Gly81, Glu111, **SO4:903**, Lys181, Lys61, **Asp146** |

Therefore the program helped reduce the complexity of virtual screening for drugs against West Nile and Dengue virus by automation of data mining of PubChem to collect data required to implement a machine learning based AutoQSAR algorithm for automatic drug lead generation. The drug leads generated by the program were required to satisfy the Lipinki’s drug-likeness criteria. Further, by feeding the structure of the drug leads generated by the program as programmatic inputs to AutoDock-Vina for *In Silico* modelling of ligand-protein interaction, the process of virtual screening by automated. Thus, the program helps achieve complete automation in identifying drugs against West Nile and Dengue virus which further has to be examined experimentally for drug potential through in *In Vitro* and *In Vivo* testing.

## Conclusion and future scope

The present work has implemented programmatic data mining of PubChem database to automate the data acquisition to implement a machine learning based AutoQSAR algorithm to generate drug leads for West Nile and Dengue virus which belong to the family of flaviviruses. The real time data fetching from PubChem is a distinguishing feature of the work which makes for a dynamic drug lead generation approach. The drug leads generated by the program were required to satisfy the Lipinski’s criteria for likelihood to possess oral bio-availability. The structure of the drug lead compounds were programmatically downloaded and fed as programmatic inputs to AutoDock-Vina by the program for an automated *In Silico* study of interaction of drug lead compounds and West Nile virus drug target identified with PDB ID : 2Oy0 and the Dengue virus drug target identified with the PDB ID : 1L9K. The results showed favourable interaction with the drug lead compounds generated by the program and the West Nile and Dengue virus drug target methyltransferase and thus making the program a useful tool in automating the process of virtual screening and identifying drugs against West Nile and Dengue virus. While the drug leads generated by the program must be further tested for drug potential through experimental testing by the experimental research community there is still scope to make the automation algorithm more technically aware of technical nuances to increase the prediction and drug identification accuracy through computational means. This our research team endeavours to pursue while also calling the attention of the computational research community to pursue the same.

## Notes

### Competing Interest Statement

The authors have declared no competing interest.

## References

1. Kim, S., Thiessen, P. A., Bolton, E. E., Chen, J., Fu, G., Gindulyte, A., Bryant, S. H. (2015). PubChem Substance and Compound databases. Nucleic Acids Research, 44(D1). doi:10.1093/nar/gkv951

2. Kim, S., Thiessen, P. A., Bolton, E. E., & Bryant, S. H. (2015). PUG-SOAP and PUG-REST: Web services for programmatic access to chemical information in PubChem. Nucleic Acids Research, 43(W1). doi:10.1093/nar/gkv396

3. Swain, M. (2014). PubChemPy: A way to interact with PubChem in Python.

4. Bhardwaj, V. (2014). Quantitative structure–activity relationship (QSAR) studies as strategic approach in drug discovery. Medicinal Chemistry Research, 23(12), 4991–5007. doi:10.1007/s00044-014-1072-3

5. Eriksson, L., & Johansson, E. (1996). Multivariate design and modeling in QSAR. Chemometrics and Intelligent Laboratory Systems, 34(1), 1–19. doi:10.1016/0169-7439(96)00023-8.

6. Bajot, F. (2009). The Use of Qsar and Computational Methods in Drug Design. Challenges and Advances in Computational Chemistry and Physics Recent Advances in QSAR Studies, 261–282. doi:10.1007/978-1-4020-9783-6_9

7. Sippl, W. (2009). 3D-QSAR – Applications, Recent Advances, and Limitations. Challenges and Advances in Computational Chemistry and Physics Recent Advances in QSAR Studies, 103–125. doi:10.1007/978-1-4020-9783-6_4.

8. Sampath, A., & Padmanabhan, R. (2009). Molecular targets for flavivirus drug discovery. Antiviral research, 81(1), 6–15.

9. Liu, L., Dong, H., Chen, H., Zhang, J., Ling, H., Li, Z., … & Li, H. (2010). Flavivirus RNA cap methyltransferase: structure, function, and inhibition. Frontiers in biology, 5(4), 286–303.

10. Fischer, A., Sellner, M., Neranjan, S., Smieško, M., & Lill, M. A. (2020). Potential Inhibitors for Novel Coronavirus Protease Identified by Virtual Screening of 606 Million Compounds. International Journal of Molecular Sciences, 21(10), 3626.

11. Rifaioglu, A., Sinoplu, E., Atalay, V., Martin, M., Cetin-Atalay, R., & Dogan, T. (2020). DEEPScreen: High Performance Drug-Target Interaction Prediction with Convolutional Neural Networks Using 2-D Structural Compound Representations. Chemical Science.

12. Gentile, F., Agrawal, V., Hsing, M., Ton, A. T., Ban, F., Norinder, U., … & Cherkasov, A. (2020). Deep Docking: A Deep Learning Platform for Augmentation of Structure Based Drug Discovery. ACS Central Science.

13. Liao, Z., You, R., Huang, X., Yao, X., Huang, T., & Zhu, S. (2019, November). DeepDock: Enhancing Ligand-protein Interaction Prediction by a Combination of Ligand and Structure Information. In 2019 IEEE International Conference on Bioinformatics and Biomedicine (BIBM) (pp. 311–317). IEEE.

14. Ton, A. T., Gentile, F., Hsing, M., Ban, F., & Cherkasov, A. (2020). Rapid identification of potential inhibitors of SARS CoV 2 main protease by deep docking of 1.3 billion compounds. Molecular informatics.

15. Mitchell, R. (2018). Web Scraping with Python: Collecting More Data from the Modern Web. O’Reilly Media, Incorporated.

16. Vanden Broucke, S., & Baesens, B. (2018). Practical Web scraping for data science (pp. 3–5). New York, NY: Apress.

17. Dixon, S. L., Duan, J., Smith, E., Von Bargen, C. D., Sherman, W., & Repasky, M. P. (2016). AutoQSAR: an automated machine learning tool for best-practice quantitative structure–activity relationship modeling. Future medicinal chemistry, 8(15), 1825–1839.

18. Kim, S., & Cho, K. H. (2019). PyQSAR: A Fast QSAR Modeling Platform Using Machine Learning and Jupyter Notebook. Bulletin of the Korean Chemical Society, 40(1), 39–44.

19. Rodgers, S. L., Davis, A. M., Tomkinson, N. P., & van de Waterbeemd, H. (2011). Predictivity of simulated ADME AutoQSAR models over time. Molecular informatics, *30*(2 3), 256–266.

20. Lipinski, C. A. (2004). Lead-and drug-like compounds: the rule-of-five revolution. Drug Discovery Today: Technologies, 1(4), 337–341.

21. Deshpande, N., Addess, K. J., Bluhm, W. F., Merino-Ott, J. C., Townsend-Merino, W., Zhang, Q., … & Kramer Green, R. (2005). The RCSB Protein Data Bank: a redesigned query system and relational database based on the mmCIF schema. Nucleic acids research, 33(suppl_1), D233–D237.

22. Jaghoori, M. M., Bleijlevens, B., & Olabarriaga, S. D. (2016). 1001 Ways to run AutoDock Vina for virtual screening. Journal of computer-aided molecular design, 30(3), 237–249.

23. Forli, S., Huey, R., Pique, M. E., Sanner, M. F., Goodsell, D. S., & Olson, A. J. (2016). Computational protein–ligand docking and virtual drug screening with the AutoDock suite. Nature protocols, 11(5), 905.

24. Tibaut, T., Borišek, J., Novič, M., & Turk, D. (2016). Comparison of in silico tools for binding site prediction applied for structure-based design of autolysin inhibitors. SAR and QSAR in Environmental Research, 27(7), 573–587.

